# Cortico-striato-pallido-thalamic loop: Effects of age on white matter neurite microstructural properties and its spatial gradients

**DOI:** 10.64898/2026.08.19.745815

**Authors:** Ekarin E. Pongpipat, Kristen M. Kennedy, Karen M. Rodrigue

## Abstract

*In-vivo* examination of neurites to understand microstructural properties of white matter tissue utilizing neurite orientation dispersion and density imaging (NODDI) has shown sensitivity to healthy aging as well as disease biomarkers and status. Neurite density index (NDI), which is a proxy for the amount of neurites, in white matter tissue typically decreases with age. However, orientation dispersion index (ODI), which is a proxy for neurite dispersion or fanning, has been mixed with studies finding both increases and decreases with age. Furthermore, white matter tracts are not uniform and hold its own unique spatial pattern or gradient in microstructural properties. In addition to the spatial pattern of the microstructural property, age-related effects have also shown spatial patterns with stronger age effects in the medial, anterior, and dorsal portions of white matter tissue. However, spatial gradients along cardinal axes within an individual’s tract have yet to be examined with age in an adult lifespan sample. The current aim of the study was to examine whether average and spatial gradients of neurite microstructural properties within tracts related to the cortico-striato-pallido-thalamic (CSPT) loop were age-sensitive. An adult lifespan sample aged 20-90 years old was recruited from the Dallas-Fort Worth metroplex (N = 104, 62% females) as part of the Dallas Area Longitudinal Lifespan Area Study (DALLAS). Participants completed an MRI session that included a structural T1-weighted image as well as multi-shell diffusion weighted imaging (MS-DWI). MS-DWI were preprocessed and tracts of interest related to the CSPT loop were obtained using probabilistic tractography. For most tracts, a significant inverted-U association with age was found for both average NDI and ODI. Most tracts revealed a reliable spatial gradient of NDI and ODI in the medial-to-lateral, posterior-to-anterior, and ventral-to-dorsal direction. Tracts related to CSPT loop were age-sensitive such that the spatial gradient was becoming more homogenous with age. This loss of spatial gradients with age is analogous to network-level dedifferentiation observed in BOLD functional connectivity. These findings highlight that age effects in a fundamental circuit for both basic and higher-order function is significantly age sensitive and while organized into spatial gradients, these gradients are also vulnerable to aging.

## INTRODUCTION

Reduction in white matter tissue volume with age is well-documented (Jernigan et al., 2001; Ge et al., 2002; Walhovd et al., 2005). This volume reduction is likely related to the loss of neurite integrity to the fiber bundles that comprise this tissue compartment. Beyond white matter tissue volume, studies have demonstrated that various MRI-quantifiable microstructural properties within the white matter fiber bundles are also detrimentally altered with age. White matter microstructural components such as myelin sheath and neurites are posited to decrease in integrity with age due to a cascade of cellular processes that accompany aging including a cyclic process of neuroinflammation induced cellular accumulation of reactive oxygen species (or free-radicals) that drive cellular energy deficiency (Raz and Daugherty, 2018). In animal studies, morphological alterations to the white matter fibers have been established and include dysmyelination, disorganization of axon bundles, and complex alterations to the myelin sheath such as ballooning (Peters, 2003). In human studies, the use of diffusion-weighted imaging has allowed for the estimation and examination of microstructural properties within white matter tissue at the mesoscale resolution. Diffusion tensor imaging (DTI), a single compartment model, has provided the bulk of the literature (Basser et al., 1994). Specifically, with increasing age, fractional anisotropy (FA) decreases and average speed of diffusivity increases (usually nonlinearly) in white matter bundles across the whole brain. However, this white matter fiber aging is differential, with association tracts more strongly affected than commissural and projection fibers, including the anterior thalamic radiations (ATR), cingulum bundle, corticostriatal tracts, cortico-striatal-thalamic tract, frontostriatal tract, posterior thalamic radiations (PTR), and superior longitudinal fasciculus (SLF) (Cox et al., 2016; de Groot et al., 2015; Pfefferbaum et al., 2005; Samanez-Larkin et al., 2012; Webb et al., 2020). Even among the association fiber bundles there is a pattern of differential aging noted that roughly follows a last-in, first-out, or retrogenesis order of phylogenetic and ontogenetic development (Raz, 2000; Kennedy & Raz, 2009a; Salat et al., 2005). Parallelling this, the aging literature also finds that gray matter follows a similar developmental and aging pattern, and these most age-vulnerable regions are those that support higher-order fluid cognition that is the earliest affected in the aging process (Kennedy & Raz, 2009b; Madden et al., 2009). Thus, a focus on these vulnerable association white matter tracts stands to enable a better understanding of early, typical cognitive and brain aging. More recently, multi-shell/compartment diffusion imaging combined with biophysical modeling approaches, such as neurite orientation dispersion and density imaging (NODDI) has allowed for the more specific examination of microstructural properties of neurites (i.e., axons and dendrites). NODDI models decompose the diffusion weighted signal into 3-compartments of isotropic, intra-neurites, and extra-neurite volume fractions for each voxel. Using biologically-informed rates of diffusion as fixed parameters for isotropic diffusivity as well as parallel and perpendicular diffusivity for intra- and extra-neurites compartments, NODDI also has one less estimated parameter than DTI. Use of NODDI overcomes tensor-based single-shell assumptions by accounting for the different compartments of the diffusion weighted signal rather than a single compartment and uses biologically-informed parameters to reduce the number of estimations producing more precise estimation of the neurite density index (NDI), orientation dispersion index (ODI), and free water fraction (FWF) (Zhang et al., 2012). Larger NDI values represent higher amounts of neurite or axons in white matter tissue and larger ODI values represent higher dispersion, fanning or axonal complexity. In the aging literature using these metrics, both indices have been shown to be altered with age in white matter tissue, NDI fairly consistently, but mixed associations have been found in ODI. Specifically, with increasing age, NDI decreases in most association fiber bundles, including the ATR, cingulum bundle, PTR, and SLF (Cox et al., 2016; Raghavan et al., 2021; Bauer et al., 2022). ODI has been reported to increase in the cingulum with age, but decrease in ATR and PTR (Cox et al., 2016; Raghavan et al., 2021) and in the SLF, ODI has been reported to both increase and decrease with increasing age (Billiet et al., 2015; Cox et al., 2016; Kodiweera et al., 2016; Raghavan et al., 2021; Bauer et al., 2022). These mixed age-associations with neurite microstructural properties warrant additional studies in the healthy adult lifespan population to help clarify this pattern.

In addition to studies examining age-dependance of average white matter microstructural properties across the tissue, studies have also examined age-related spatial patterns within the white matter tissue. Along the cardinal axes (medial-to-lateral, posterior-to-anterior, and ventral-to-dorsal), stronger age-related associations with DTI microstructure (i.e., lower FA and higher MD with increasing age) are generally found in the medial, anterior, and dorsal portions of the whole white-matter tissue (Davis et al., 2009; Barrick et al., 2010; Hoagey et al., 2019) and for the anterior compared to posterior portions of the corpus collosum and uncinate fasciculus (Pfefferbaum et al., 2005; Davis et al., 2009). Posterior-to-anterior age gradients have also been reported among the segments of the corpus callosum with stronger age effects on FA and MD in anterior portions (Lebel et al. 2010; Kraft et al., 2024). Even more fine-grained analyses have found age-related associations with DTI microstructure along the tract length within several major white matter bundles, demonstrating unique patterns that peak at different locations dependent upon the tract (Davis et al., 2009; Yeatman et al., 2012; Shirazi et al., 2021). NODDI microstructural properties along the tract length of major white matter bundles have also shown developmental associations from infancy to adolescence with similarly unique patterns dependent upon the white matter tract (Lynch et al., 2020). These prior studies have examined age associations with white matter microstructure at each point along either the cardinal axes or length of a specific tract. However, we posit that microstructural properties along the cardinal axes are unique and informative to each tract and each individual. This tract- and individual-specific pattern of microstructural properties can be captured as a spatial gradient along the cardinal axes. To the best of our knowledge, no prior studies have examined individually estimated spatial gradients of neurite microstructural properties in an adult lifespan sample and warrants exploration.

The present study is particularly interested in aging of the white matter properties of the cortico-striato-pallido-thalamic (CSPT) loop as it is a foundational pathway for processing and integrating motor and cognitive information across several neuromodulatory systems (Alexander et al., 1986; Taylor & Taylor, 2000). The structural connections of this CSPT loop are highly organized and structured between regions. For example, the striatum is structurally connected with the cortex in a manner that roughly follows a posterior-to-anterior gradient. The anterior portions of striatum are connected to the anterior portions of the cortex, while the posterior portions are mostly connected to the primary sensory and motor cortices (Verstynen et al., 20120; Jarbo & Verstynen, 2015). Similarly, the mediodorsal thalamic nucleus and the prefrontal cortex show a medial-to-lateral gradient of connections. The more medial magnocellular subdivision is connected to the medial portion of the OFC and the lateral subdivision is connected to the lateral middle- and superior-frontal gyri. This structural organization can in part be explained by the striatal subcompartments of the striosome and matrix as certain tracts from the thalamic subdivisions appear to be segregated by striosome- or matrix-like compartments (Waugh et al. 2022; Funk et al., 2023). The structural organization is also explained in-part by coupled genetic expression that attracts and repels neuronal axons (Dickson et al, 2013; Fornito et al., 2019). Given the importance of this CSTP loop in various fundamental processes and its highly organized structure, the current study aimed to examine how neurite microstructural properties of white matter tracts within the CSTP loop, including its spatial gradient, may be age-sensitive.

The three main objectives in examining neurite microstructural properties within tracts related to the cortico-striato-pallido-thalamic (CSPT) loop in the adult lifespan were: 1) examine how average neurite density and orientation dispersion are related to cross-sectional age, 2) examine whether these individually estimated neurite microstructural properties are consistent and reliable across individuals at the group-level, and 3) examine whether meaningful spatial gradients of neurite microstructural properties along the cardinal axes are altered with increasing age. We tested these aims in the white matter tracts comprising the CSPT loop using fiber tractography within the anterior thalamic radiations, posterior thalamic radiations, frontoparietal tracts, the CSPT, and the whole tractogram.

## 2. METHODS

### 2.1. Participants

Participants were recruited as part of the Dallas Area Longitudinal Lifespan Aging Study (DALLAS), an adult lifespan sample from the Dallas-Fort Worth metroplex. All participants were required to be right-handed, native English speakers (i.e., acquired by age 6), that earned a high school diploma or equivalent, and had corrected-to or normal 20/20 vision and hearing (as tested in the lab). Participants were screened against and excluded if they endorsed any of the following: metabolic or neurological disorders, traumatic brain injury with loss of consciousness > 5 min, psychiatric disorders, history of present substance abuse, psychoactive or cognition-altering medications, cardiovascular and metabolic diseases, diabetes, and for ferrous metal or claustrophobia that would interfere with obtaining an MRI scan. Participants were also screened against signs of depression (excluded if Center for Epidemiologic Studies Depression Scale (CES-D) score > 16 out of 20; Radloff, 1977) or dementia (excluded if Mini-Mental State Examination score < 25 out of 30; Folstein et al., 1975). All participants provided written signed informed consent that was approved by both the University of Texas at Dallas and the University of Texas at Southwestern institutional review boards before participating in the study. The present sample consisted of participants from the second wave of the DALLAS study data collection as that is when multi-shell diffusion-weighted imaging was introduced. Participants included 104 individuals aged 20-90 years of age (65 females). See **Table 1** for sample demographics. Age was evenly distributed between men and women and was unrelated to years of education and depression score but negatively correlated with MMSE score.

**Table 1.** Participant Demographics.

| Variable | Adult Age Group |  |  |  | Overall<br>(20 - 90) | Statistic | p-value |
| --- | --- | --- | --- | --- | --- | --- | --- |
|  | Younger<br>(20-39) | Middle<br>(40-59) | Older<br>(60-72) | Oldest<br>(73 - 90) |  |  |  |
| N | 26 | 26 | 25 | 27 | 104 |  |  |
| Age | 30.88<br>(4.89) | 51.81<br>(5.40) | 65.72<br>(3.69) | 79.00<br>(4.52) | 57.09<br>(18.74) |  |  |
| Women | 17<br>(65.38%) | 14<br>(53.85%) | 13<br>(52.00%) | 20<br>(76.92%) | 65<br>(62.50%) | $\chi^2(1) = 0.49$ | 0.622 |
| Education<br>(Years) | 16.38<br>(1.70) | 15.31<br>(2.46) | 16.08<br>(2.41) | 16.00<br>(2.71) | 15.94<br>(2.34) | $t(102) = -0.13$ | 0.894 |
| CESD | 5.67<br>(5.90) | 5.08<br>(6.00) | 4.92<br>(6.75) | 4.62<br>(4.46) | 5.07<br>(5.72) | $t(100) = -0.66$ | 0.512 |
| MMSE | 29.60<br>(0.65) | 28.85<br>(1.16) | 29.16<br>(0.69) | 28.35<br>(1.16) | 28.95<br>(1.08) | $t(101) = -4.18$ | < .001 |

*Note:* Means and (standard deviations) are reported for baseline age, years of education, CESD, and MMSE. Age categories are only reported for descriptive purposes as age is treated as a continuous variable in all analyses. There are no significant effects of age on sex, years of education, CESD, but MMSE decreased significantly with increasing age. Abbreviations: CESD - Center for Epidemiological Studies – Depression, MMSE - Mini-Mental State Examination.

### 2.2. MRI Acquisition

Several multi-modal MRI sequences were collected at the University of Texas Southwestern Medical Center’s Advanced Imaging Research Center on a 3 Tesla Philips Achieva Scanner with a 32-channel head coil using SENSE acceleration, including the following sequences: 1. An anatomical T1-weighted (T1w) image using the Magnetization Prepared Rapid Gradient Echo (MP-RAGE) sequence in the sagittal direction with an isotropic voxel size of 1mm^3^ (matrix: 170 [Left-Right; LR] x 240 [Posterior-Anterior; PA] x 256 [Inferior-Superior; IS]; field of view [FOV]: 170mm [LR] x 240mm [PA] x 256mm [IS]; repetition time [TR]: 8.13ms; echo time [TE]: 3.73ms; flip angle: 12°; acquisition time: 4:12), 2. A multi-shell diffusion-weighted image (DWI-MS) using an Echo Planar Imaging (EPI) sequence in the axial direction with a reconstructed voxel size of 1.96mm x 1.96mm x 2.2mm (matrix: 100 [LR] x 99 [PA]; FOV: 220mm [LR] x 220mm [PA] x 158mm [IS]; TR: shortest (6431 ± 7.78ms); TE: 105.74ms; flip angle: 90°; acquired voxel size: 2.2mm [LR] x 2.2mm [PA] x 2.2mm [IS]; multiband factor = 2; acquisition time: 13min). The multi-shell DWI series contained 16 b = 0 s/mm^2^, 30 b = 1000 s/mm^2^, 30 b = 2500 s/mm^2^, and 30 b = 4000 s/mm^2^ gradient volumes. 3. Two field map images with identical geometry to the DWI images were also collected to correct for susceptibility-induced and eddy-current distortions. The field maps images consisted of two non-diffusion gradient volumes (b = 0 s/mm^2^) with opposing phase encoding directions (one volume from anterior-to-posterior and another volume from posterior-to-anterior).

### 2.3. MRI Preprocessing

T1w images were preprocessed to obtain regions of interest (ROIs) and avoidance (ROAs) for the generation of tracts related to the cortical-striato-pallido-thalamic-cortical loop. These regions were parcellated at the individual level rather than using a pre-defined atlas to allow for more individual-specificity (see Figure 1A-B). T1w images were initially processed using *Freesurfer’s recon-all* to obtain cortical ROIs and ROAs (v5.3; Dale et al., 1999). The output was visually inspected and manual modifications were performed, as needed, to ensure the appropriate formation of pial and white matter boundaries. The cortex was parcellated using the Desikan-Killiany atlas as well as the Brodmann’s atlas into frontal and parietal aggregate regions (Dale et al., 1999; Desikan et al., 2006; Pijnenburg et al., 2021). The frontal cortex mask consisted of the rostral anterior cingulate, caudal anterior cingulate, superior frontal, rostral middle frontal, and caudal middle frontal gyri as well as the pars orbitalis, pars triangularis, and pars opercularis gyri. Brodmann area 6 was used as an exclusionary mask for the frontal cortex to avoid including supplemental motor areas. The parietal ROI consisted of the supramarginal gyrus, inferior parietal/angular gyrus (i.e., Brodmann area 39), superior parietal lobule, precuneus, and isthmus cingulate. The posterior cingulate was not included since the region lies more central underneath motor-associated areas. The occipital lobe portion of the inferior parietal ROI was removed by using occipital regions as defined by Brodmann’s areas 17, 18, and 19 in a subtraction mask. Subcortical regions were generated using *FSL’s FIRST* as well as *HIPS-THOMAS*. Specifically, the basal ganglia were parcellated into the caudate, putamen, nucleus accumbens, and pallidum using *FSL’s FIRST* (Patenaude et al., 2011) and the thalamus was parcellated using *Histogram-based Polynomial Synthesis - Thalamus Optimized Multi-Atlas Segmentation* (*HIPS-THOMAS*; Su et al., 2019).

**Figure 1.**
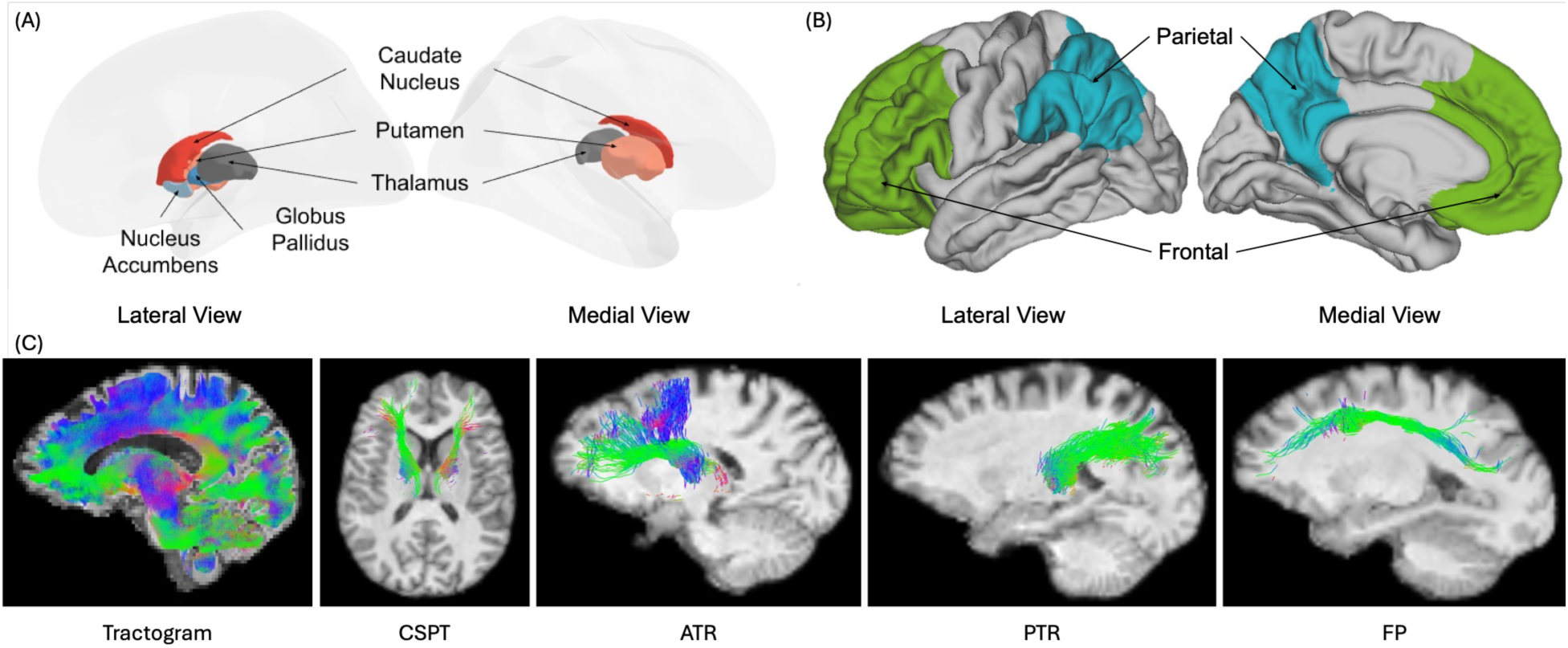
Illustration of tracts of interest generated using probabilistic tractography guided by specified regions of interest. Subcortical and cortical regions of interest and avoidance (A-B). Tracts of interest are shown for a representative subject (C). Abbreviations: ATR - anterior thalamic radiation, F – frontal, FP - fronto-parietal, CSPT - cortico-striatal-pallido-thalamic, P – parietal, PTR - posterior thalamic radiation

DWI-MS data were preprocessed using *FSL v6.0.5.2* (Andersson et al., 2003; Smith et al., 2004) to correct for susceptibility-induced distortions (*topup*; Andersson et al., 2003; Smith et al., 2004) and eddy-induced distortions along with motion (*eddy*; Andersson et al., 2016a; Andersson et al., 2016b). The brain and regional masks were warped into the individual’s preprocessed DWI-space by warping the T1w extracted brain to the mean b = 0 s/mm^2^ using *FSL’s flirt* with boundary-based registration and 6 degrees of freedom (rotation and translation along the x, y, z axes) while respecting the white and gray matter interface (Greve et al., 2009; Jenkinson et al., 2001; Jenkinson et al., 2002). Neurite orientation dispersion and density imaging (NODDI) with the Watson distribution was modeled and reconstructed using the *Accelerated Microstructure Imaging via Convex Optimization* (*AMICO*) software (Daducci et a., 2015; Zhang et al. 2012). For each voxel, the reconstructed NODDI model provided estimates of free water (isotropic) volume fraction (FWF), neurite density index (NDI), and orientation dispersion index (ODI). Tissue weighting was applied to NDI and ODI by first subtracting the FWF from 1 to obtain a tissue volume fraction image (TVF) and then multiplying that by NDI and ODI, respectively (Parker et al., 2021). These tissue-weighted NDI and ODI metrics were the indices of interest used in subsequent analyses.

### 2.4. Tractography

Probabilistic tractography was performed using the *MRtrix3* software (Tournier et al., 2019). Data from the T1w image were classified into one of five tissue types (i.e., cortical gray matter, subcortical gray matter, white matter, cerebral spinal fluid, and pathological tissue such as tumors) based on a hybrid surface and volume segmentation (HSVS) using both the *Freesurfer* and *FSL* software, respectively (*5ttgen*; Smith et al., 2020). Multi-shell, multi-tissue constrained spherical deconvolution (MSMT-CSD) reconstruction was performed to estimate the response function and orientation distribution function (ODF) for cerebral spinal fluid, gray matter tissue, and white matter tissue (*dwi2response msmt_5tt* and *dwi2fod msmt_csd*; Jeurissen et al., 2014). Utilizing information from the MSMT-CSD, probabilistic tractography was performed using the 2nd order integration over fiber orientation distributions (iFOD2) algorithm (*tckgen iFOD2*; Tournier et al., 2010; 2012). White matter tracts were seeded using the gray matter-white matter interface created from the five-tissue type classification and constrained to anatomical tracts via the Anatomically Constrained Tractography algorithm (Smith et al., 2020; Smith et al., 2012). Each white matter tract was seeded up to 1 million iterations at the gray-white matter interface and terminated early if 10,000 streamlines followed its respective regions of interest and avoidance. Similarly, the whole brain tractogram was seeded up to 1 million streamlines at the gray-white matter interface but did not terminate early.

### 2.5. Tracts of Interest (TOIs)

Tracts of interest (TOIs) included the cortico-striato-pallido-thalamic (CSPT) tracts, anterior thalamic radiations (ATR) tracts, posterior thalamic radiations (PTR) tracts, and the fronto-parietal (FP) tracts as well as the whole brain tractogram. Each TOI was generated using tract-specific angles as well as regions of interest and avoidance (see **Table 2** and **Figure 1C**). For each tract of interest, streamlines were required to pass through each ROI in the specified ROI order. The reversed direction was also performed, and both directions were combined to form a single tract bundle. Except for the whole brain tractogram, each tract bundle was estimated separately for each hemisphere. For average microstructural property, each white matter TOI was converted to its own masked volume and the mean value was extracted. For the whole brain tractogram, left and right hemispheric masks of the T1w in MNI space were warped to DWI space to create left and right hemispheres of the tractogram. Microstructural properties were combined across hemispheres if their values had a correlation of 0.7 or higher.

**Table 2.**
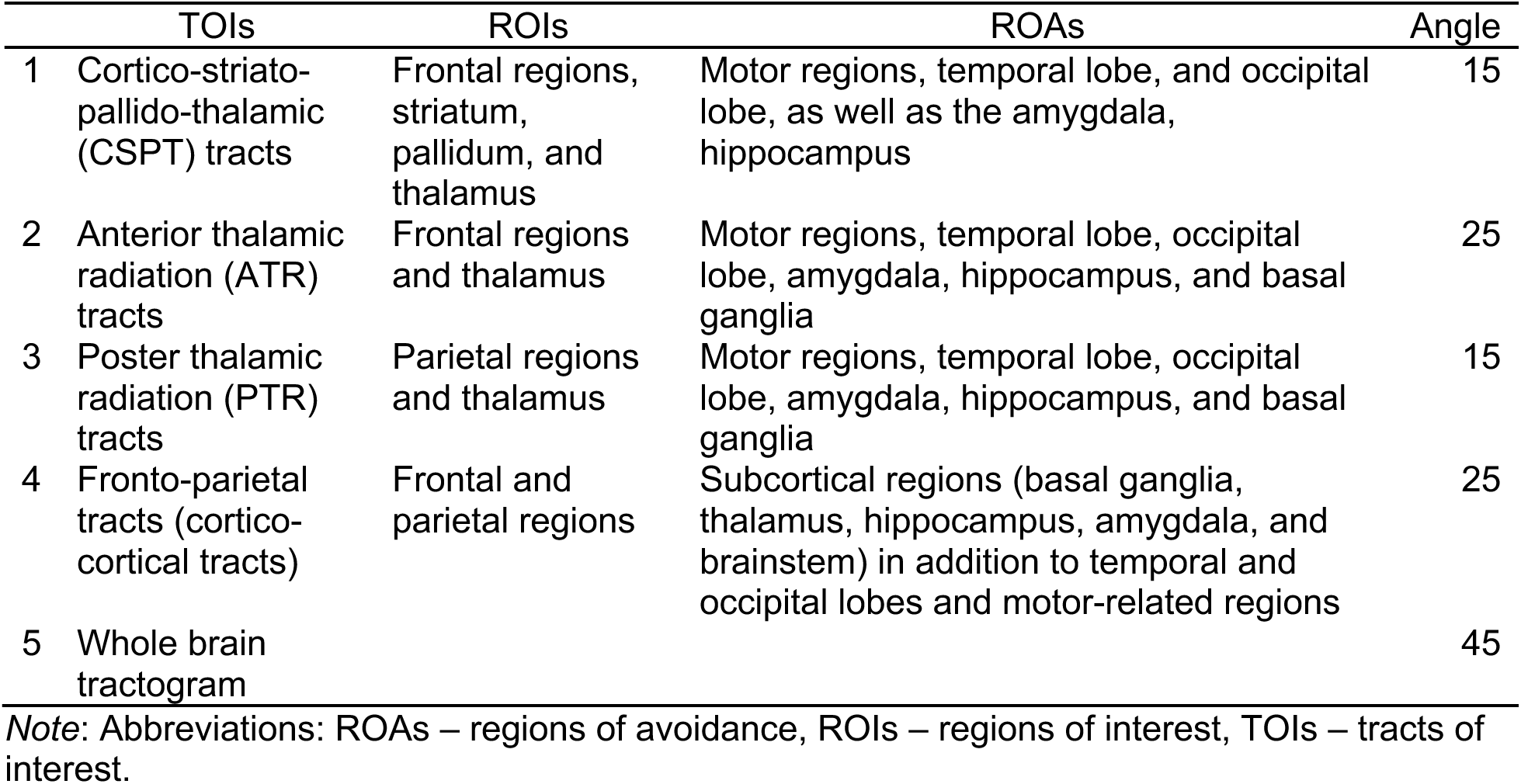
Specified parameters for each tract of interest.

### 2.6. Individual-Level Spatial Gradients

For spatial gradients, the masked volume file of each white matter TOI was simultaneously warped to MNI space via the T1w scan using ANTs (Tustison et al., 2021). For each TOI, the coordinates as well as their respective NODDI microstructural properties were extracted. Outlier values of the microstructural property from the entire mask were removed based on robust z-score to avoid incorporating values that might contain noise or information about other tissue compartments (Leys et al., 2013; Theriault et al., 2024). These coordinates and neurite microstructural properties were used to estimate the spatial gradient along the medial-to-lateral, anterior-to-posterior, and ventral-to-dorsal directions. Spatial gradients were estimated for each individual by running a general linear model (GLM) with the microstructural property as the dependent variable and the location along the medial-to-lateral, posterior-to-anterior, and ventral-to-dorsal axes as the variables of interest for each individual, tract of interest (TOI), and NODDI metric (see **Figure 2** for pipeline visualization).

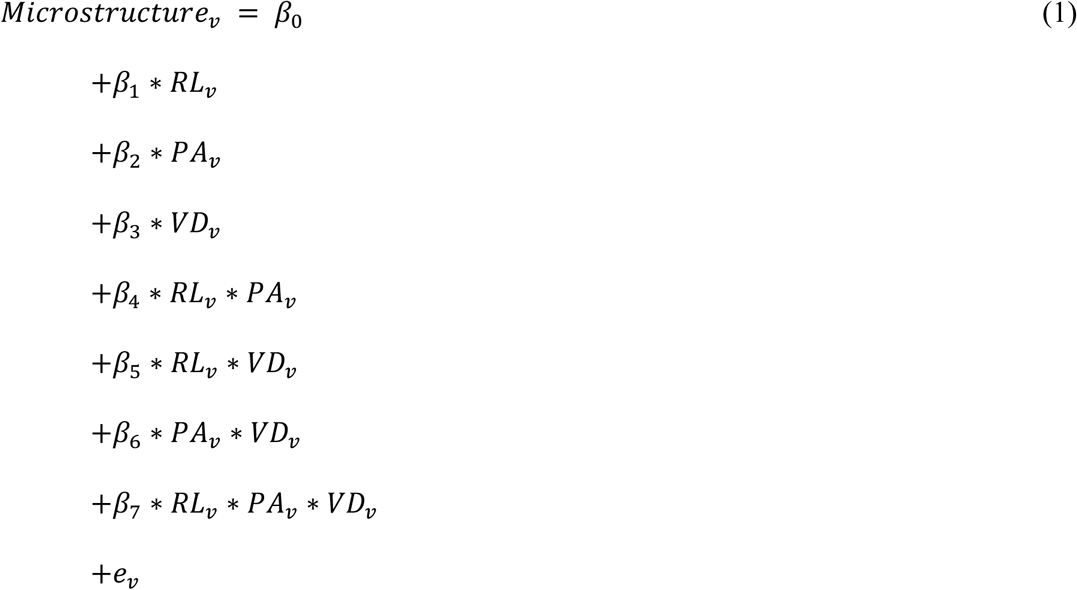

**Figure 2.**
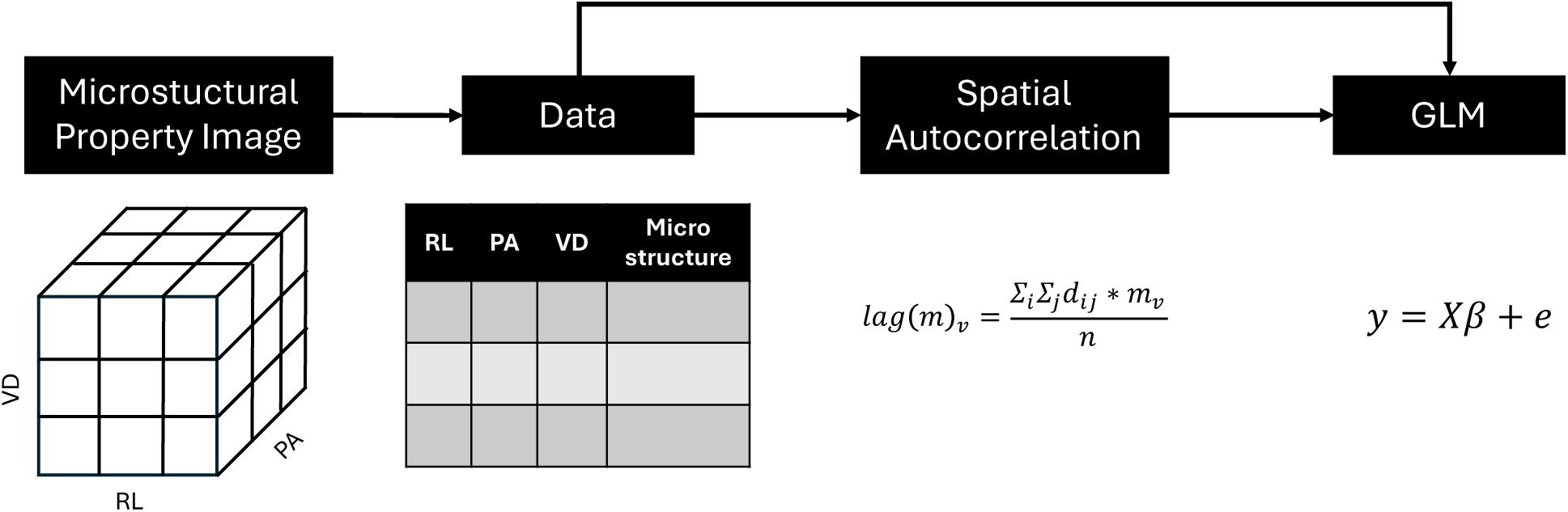
Pipeline for spatial gradient extraction and analysis for each tract of interest at the individual level. Abbreviations: VD – ventral-dorsal, RL – right-left, PA – posterior-anterior

where *v* represents each voxel within a TOI, RL represents the right-left axis that is interpreted as medial-lateral dependent upon the sign of the estimate and the hemisphere of the TOI (i.e., the sign was flipped for TOIs from the right hemisphere), PA represents the posterior-anterior axis, and VD represents the ventral-dorsal axis. Although a more complex GLM that incorporates nonlinearity (e.g., quadratic and cubic functions) would allow for more precision of the spatial gradients, the interpretation becomes increasingly complex. Thus, the current spatial gradient model balances complexity and interpretability by allowing each spatial axis to be estimated simultaneously while still allowing the estimates to be readily interpretable. The 3-way interaction of each dimension allowed the lower-order effects to be interpreted as on average across the other dimensions rather than controlling for the other dimensions. The spatial gradient estimate in each cardinal direction (*β*_1_, *β*_2_, and *β*_3_) of only the lower-order terms was then taken and analyzed at the group level.

To ensure that the results were not merely due to spatial autocorrelation, a spatially lagged variable as well as Moran’s index was calculated. The spatially lagged microstructural property, *lag*(*m*)*_v_*, which is conceptually similar to a temporally lagged variable, was calculated by creating a distance matrix of each voxel pair within an ROI, multiplying each distance by its respective voxel’s microstructural property, and then taking the mean of each voxel. Only voxels that were either touching at the sides, edges, or corners were used to obtain the spatial lag of the nearest neighboring voxel.

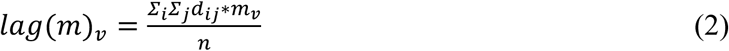

The spatially lagged microstructural property for its respective microstructural property was used as a nuisance variable at the individual level:

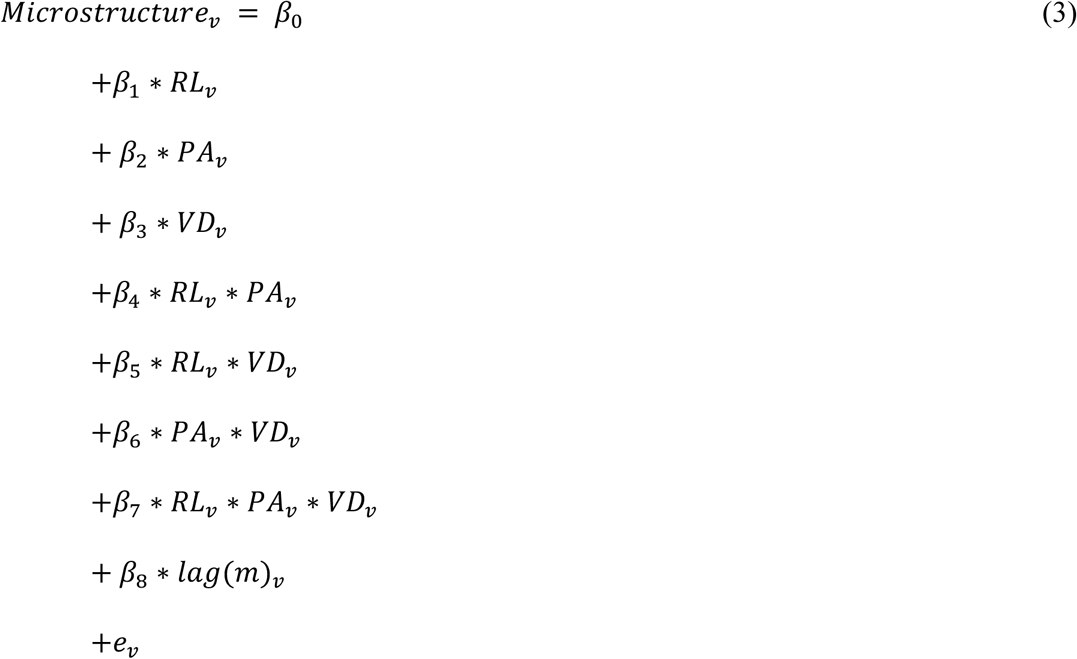

In addition to the spatially lagged variable, spatial autocorrelation was examined using Moran’s index (Moran, 1948; Chen, 2013). Moran’s index (*I*), in matrix notation, is 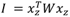, where *x_z_* represents the z-scored microstructural property, 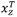 represents the transposed version of *x_z_*, and W represents a distance matrix of each voxel pair normalized such that the sum of the W matrix sums to 1. The algebraically expanded version of Moran’s index is essentially a modified correlation, which is conceptually similar to the temporal autocorrelation. Moran’s index was used as a nuisance variable at the group level. Respective of cardinal direction and microstructural property, spatial gradient estimates were combined across hemispheres if their values were correlated at 0.7 or greater.

### 2.7. Group-Level Analyses

To understand the age sensitivity and trajectories of NODDI microstructural properties, general linear models (GLMs) were performed. For mean neurite microstructural property, the dependent variable (DV) was either mean NDI or mean ODI with linear and quadratic age as the independent variables (IV) while controlling for sex. If there was no significant effect involving quadratic age for a particular TOI, the quadratic age term was removed from the model and reanalyzed with only linear age. Given the number of non-independent analyses, multiple comparison correction was performed using the *M_eff_* method (Cheverud, 2001; Nyhold, 2004; Derringer, 2018). *M_eff_* was calculated separately for each neurite microstructural property. For these analyses, the significant *α_Meff_* threshold was 0.023 (*M_eff_* = 2.20) for mean NDI and 0.010 (*M_eff_* = 4.91) for mean ODI. Exploratory analyses were additionally performed to determine whether the mean amount of microstructural property as well as its age trajectory differed from the whole brain tractogram. These exploratory analyses were performed using a linear mixed effects model with Satterthwaite denominator degrees of freedom (Kuznetsova et al., 2017). The model included either mean NDI or mean ODI as the DV with the IVs being the tracts of interest, linear age, and quadratic age along with its interactions while controlling for sex and allowing for a random intercept for each subject. The tracts of interest were dummy-coded with the whole brain tractogram serving as the reference level.

For analyses involving spatial gradients, individually estimated gradients were first examined to ensure that the gradients existed and were consistent at the group-level. This analysis was performed for each tract of interest, microstructural property, and cardinal direction by using a GLM with the spatial gradient estimate as the DV and the IV being the intercept while controlling for the mean-centered Moran’s index. Significant spatial gradients were then analyzed to determine age sensitivity and trajectory using a similar GLM as analysis with mean neurite microstructural properties. Specifically, the DV was the spatial gradient estimate and the IV was linear and quadratic age while controlling for sex and Moran’s index. Similar to the mean neurite microstructural property analyses, if a model had no significant effect involving quadratic age, the term was removed and reanalyzed with only linear age. The sign was flipped for negative spatial gradient estimates to allow for the interpretation that higher values represent a steeper spatial gradient (or more microstructural property on one end of a cardinal axis relative to the other end), while lower values represent a less steep spatial gradient (or more homogenous amounts of microstructural properties across the cardinal axis). To account for the number of independent analyses, *M_eff_* was calculated separately for each microstructural property and cardinal direction. For these analyses, the significant *α_Meff_* threshold was 0.0051 for NDI_ML_ (*M_eff_* = 9.86), 0.0051 for NDI_PA_ (*M_eff_* = 9.76), 0.0058 for NDI_VD_ (*M_eff_* = 8.70), 0.0051 for ODI_ML_ (*M_eff_* = 9.87), 0.0051 for ODI_PA_ (*M_eff_* = 9.81), and 0.0052 for ODI_VD_ (*M_eff_* = 9.71). An exploratory analysis was also performed to determine how linear and quadratic age influences microstructural properties at specific slices along the cardinal directions similar to prior studies (Hoagey et al., 2019; Pfefferbaum et al., 2005). Specifically, a GLM was performed with mean microstructural property as the DV and linear and quadratic age as the IVs while controlling for sex. This analysis was performed for each microstructural property, cardinal direction, and slice along the cardinal direction. All group analyses were performed using *R v4.1.0* within the *R Studio IDE* (R Core Team, 2023; Posit team, 2024).

## 3. RESULTS

### 3.1. Age Effects on Mean Neurite Microstructural Properties

Significant quadratic age effects were observed for both mean NDI and ODI for all tracts of interest. In all TOIs the quadradic function reflects that the average neurite metric increased slightly until middle-age and subsequently decreased in older adulthood. See **Table 3** for individual model statistical coefficients and **Figure 3** for illustration of age-regression curves for each tract for NDI and ODI metrics. For mean NDI, significant quadratic effects were found for the cortico-striato-pallido-thalamic, frontoparietal, anterior thalamic radiation, and posterior thalamic radiation tracts as well as the whole brain tractogram (*p*’s < 0.011, adjusted *R^2^* ranged from 0.06 to 0.14). For mean ODI, significant quadratic age effects were found for the combined cortico-striato-pallido-thalamic, bilateral anterior thalamic radiation, combined posterior thalamic radiation, and bilateral frontoparietal tracts as well as the whole brain tractogram (*p*’s < 0.0005, adjusted *R^2^* ranged from 0.11 to 0.30).

**Figure 3.**
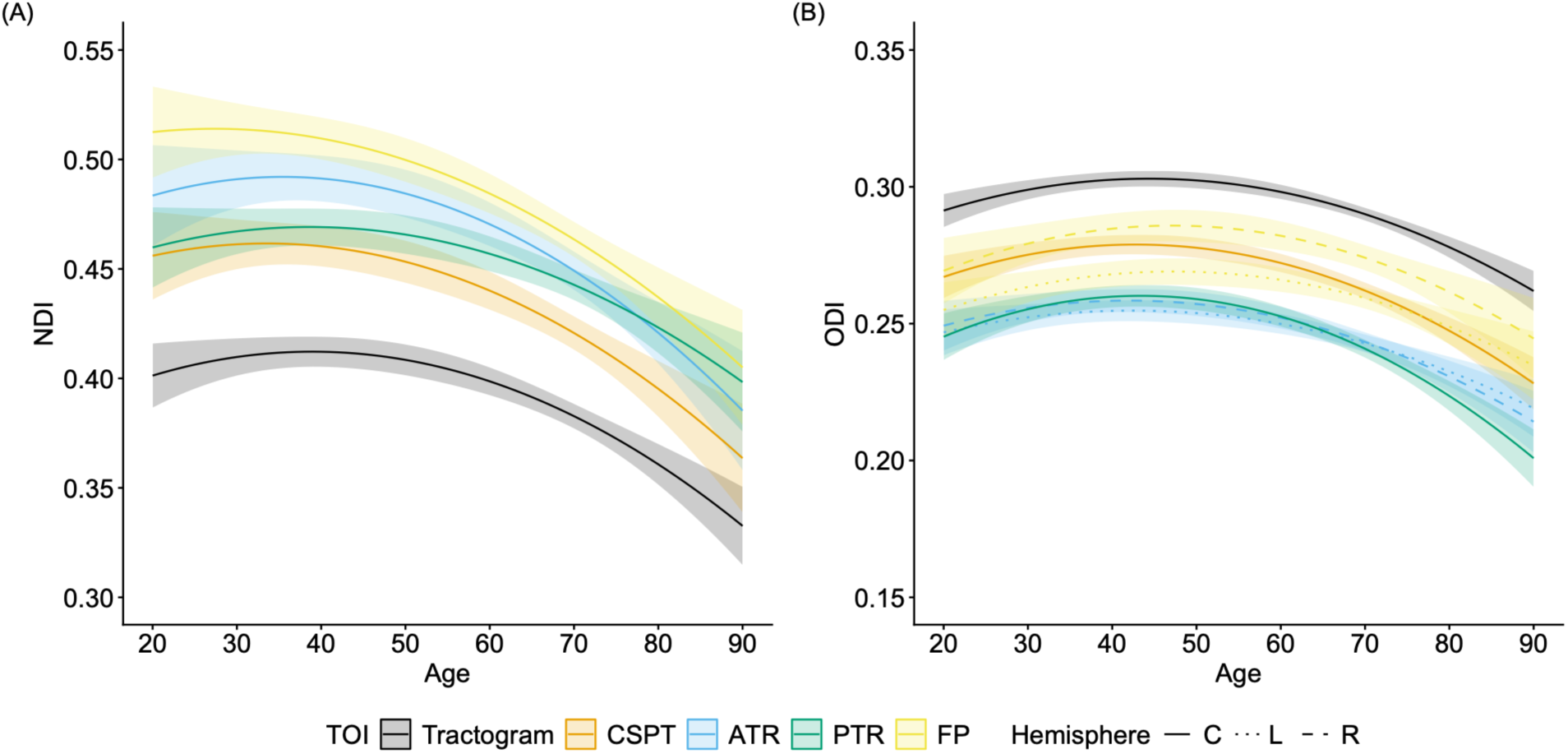
Significant quadratic age effects on average neurite density index (A) and orientation dispersion index (B) for five sets of white matter tracts. Note: The y-axis represents the estimated neurite microstructural property when controlling for sex. Bands represent the 95% confidence interval. Abbreviations: ATR – anterior thalamic radiations, C – combined, CSPT - cortico-striato-pallido-thalamic tract, FP – frontoparietal tract, L – left, NDI – neurite density index, ODI – orientation dispersion index, PTR – posterior thalamic radiations, R - right.

**Table 3.** Quadratic age effects on mean neurite density and orientation dispersion indices.

| Metric | IV | TOI | *b* | *t* | *p* | 95% CI |  | *R*<sup>2</sup><sub>Adj.</sub> |
| --- | --- | --- | --- | --- | --- | --- | --- | --- |
|  |  |  |  |  |  | LL | UL |  |
| NDI | Age<sup>2</sup> | Tractogram (C) | -3.05 ×<br>10<sup>-5</sup> | -4.162 | 6.833 ×<br>10<sup>-5</sup> | -4.51 ×<br>10<sup>-5</sup> | -1.60 ×<br>10<sup>-5</sup> | 0.143 |
|  |  | CSPT (C) | -3.07 ×<br>10<sup>-5</sup> | -2.997 | 3.485 ×<br>10<sup>-3</sup> | -5.11 ×<br>10<sup>-5</sup> | -1.04 ×<br>10<sup>-5</sup> | 0.078 |
|  |  | ATR (C) | -3.59 ×<br>10<sup>-5</sup> | -3.126 | 2.343 ×<br>10<sup>-3</sup> | -5.86 ×<br>10<sup>-5</sup> | -1.31 ×<br>10<sup>-5</sup> | 0.083 |
|  |  | PTR (C) | -2.69 ×<br>10<sup>-5</sup> | -2.862 | 5.186 ×<br>10<sup>-3</sup> | -4.55 ×<br>10<sup>-5</sup> | -8.23 ×<br>10<sup>-6</sup> | 0.070 |
|  |  | FP (C) | -2.78 ×<br>10<sup>-5</sup> | -2.581 | 1.141 ×<br>10<sup>-2</sup> | -4.92 ×<br>10<sup>-5</sup> | -6.41 ×<br>10<sup>-6</sup> | 0.057 |
| ODI | Age<sup>2</sup> | Tractogram (C) | -1.97 ×<br>10<sup>-5</sup> | -6.517 | 3.220 ×<br>10<sup>-9</sup> | -2.57 ×<br>10<sup>-5</sup> | -1.37 ×<br>10<sup>-5</sup> | 0.297 |
|  |  | CSPT (C) | -2.28 ×<br>10<sup>-5</sup> | -5.764 | 1.039 ×<br>10<sup>-7</sup> | -3.06 ×<br>10<sup>-5</sup> | -1.49 ×<br>10<sup>-5</sup> | 0.253 |
|  |  | ATR (L) | -1.58 ×<br>10<sup>-5</sup> | -3.678 | 3.941 ×<br>10<sup>-4</sup> | -2.43 ×<br>10<sup>-5</sup> | -7.26 ×<br>10<sup>-6</sup> | 0.118 |
|  |  | ATR (R) | -1.91 ×<br>10<sup>-5</sup> | -4.100 | 8.829 ×<br>10<sup>-5</sup> | -2.84 ×<br>10<sup>-5</sup> | -9.87 ×<br>10<sup>-6</sup> | 0.144 |
|  |  | PTR (C) | -2.73 ×<br>10<sup>-5</sup> | -6.258 | 1.151 ×<br>10<sup>-8</sup> | -3.60 ×<br>10<sup>-5</sup> | -1.86 ×<br>10<sup>-5</sup> | 0.287 |
|  |  | FP (L) | -1.89 ×<br>10<sup>-5</sup> | -3.633 | 4.585 ×<br>10<sup>-4</sup> | -2.92 ×<br>10<sup>-5</sup> | -8.55 ×<br>10<sup>-6</sup> | 0.115 |
|  |  | FP (R) | -2.24 ×<br>10<sup>-5</sup> | -3.654 | 4.308 ×<br>10<sup>-4</sup> | -3.46 ×<br>10<sup>-5</sup> | -1.02 ×<br>10<sup>-5</sup> | 0.118 |

*Note:* Abbreviations: ATR - anterior thalamic radiation, C – combined, CSPT - cortico-striato-pallido-thalamic, FP – frontoparietal, L – left, NDI - neurite density index, ODI - orientation dispersion index, TOI - tract of interest, PTR - posterior thalamic radiation, R - right.

For each microstructural property, an exploratory LME analysis was performed to determine how the amount of microstructural property and its age trajectory for each tract of interest potentially differed from the whole brain tractogram, which was used as a reference for overall white matter tissue (see **Table 4** and **Table 5**). For mean neurite density index, each tract of interest had higher levels of NDI compared to the whole brain tractogram (*p* < 2.783 × 10^−31^, adjusted *R^2^* ranged from 0.30 to 0.64). The whole brain tractogram showed both a significant linear age (*p* = 1.784 × 10^−8^, adjusted *R^2^* = 0.17) and quadratic age effect (*p* = 1.647 × 10^−3^, adjusted *R^2^* = 0.05), similar to the above GLM results. For the linear age trajectory, the cortico-striato-pallido-thalamic tract, anterior thalamic radiation, and frontoparietal tracts evidenced a steeper age slope compared to the whole brain tractogram (*p*’s < 0.011, ATR: *p* = 0.013, adjusted *R^2^*ranged from 0.01 to 0.05). In other words, average NDI decreased faster with age for these tracts compared to the whole brain tractogram. However, no significant differences for the quadratic age analyses were detected in any of the TOIs relative to the whole brain tractogram suggesting that the other TOIs followed a similar quadratic trajectory as general white matter.

**Table 4.**
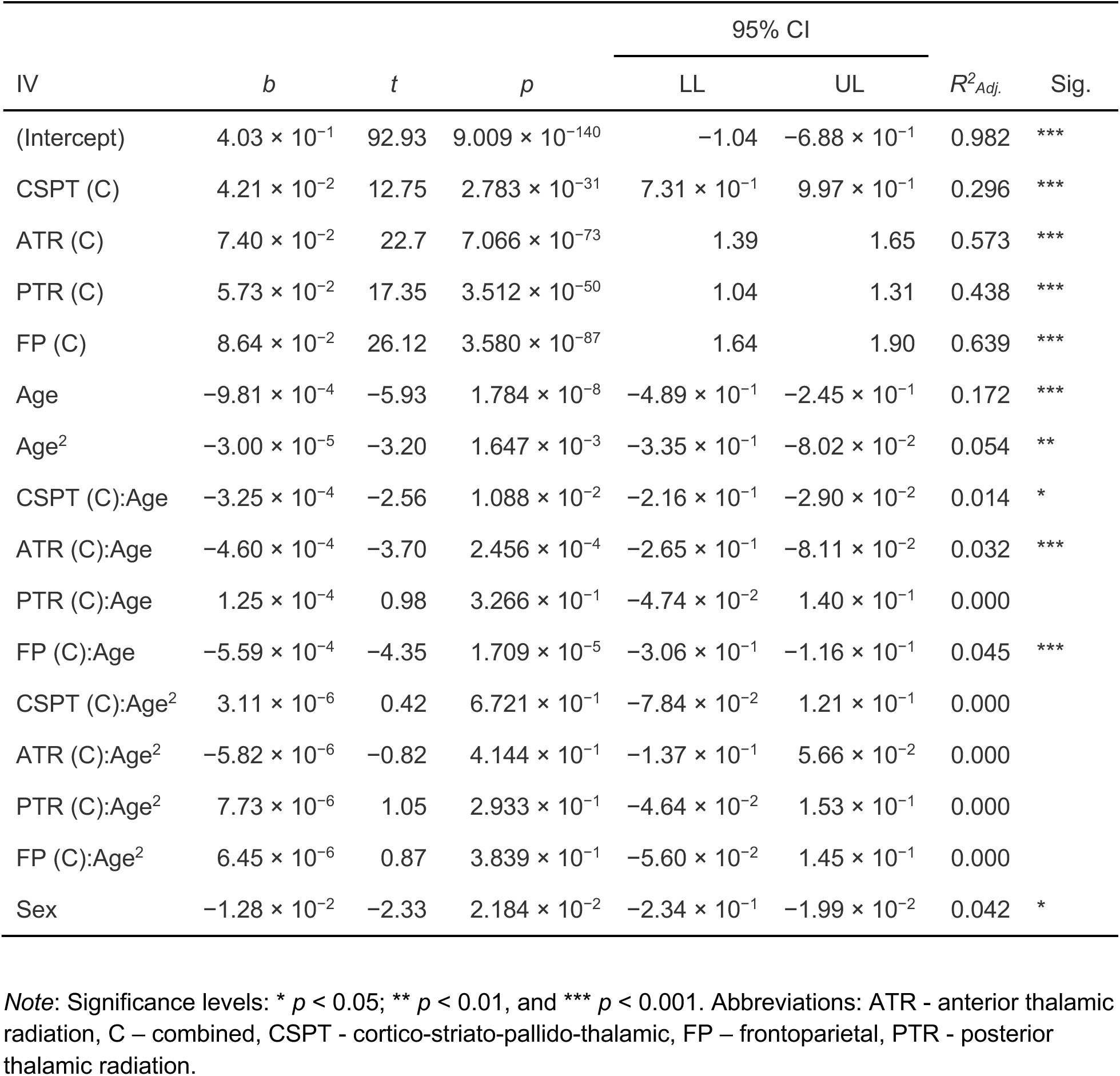
Age effect comparisons of mean neurite density index within specific tracts relative to the whole tractogram.

| IV | $b$ | $t$ | $p$ | 95% CI | | $R^2_{Adj.}$ | Sig. |
| --- | --- | --- | --- | --- | --- | --- | --- |
|  |  |  |  | LL | UL |  |  |
| (Intercept) | $4.03 \times 10^{-1}$ | 92.93 | $9.009 \times 10^{-140}$ | -1.04 | $-6.88 \times 10^{-1}$ | 0.982 | *** |
| CSPT (C) | $4.21 \times 10^{-2}$ | 12.75 | $2.783 \times 10^{-31}$ | $7.31 \times 10^{-1}$ | $9.97 \times 10^{-1}$ | 0.296 | *** |
| ATR (C) | $7.40 \times 10^{-2}$ | 22.7 | $7.066 \times 10^{-73}$ | 1.39 | 1.65 | 0.573 | *** |
| PTR (C) | $5.73 \times 10^{-2}$ | 17.35 | $3.512 \times 10^{-50}$ | 1.04 | 1.31 | 0.438 | *** |
| FP (C) | $8.64 \times 10^{-2}$ | 26.12 | $3.580 \times 10^{-87}$ | 1.64 | 1.90 | 0.639 | *** |
| Age | $-9.81 \times 10^{-4}$ | -5.93 | $1.784 \times 10^{-8}$ | $-4.89 \times 10^{-1}$ | $-2.45 \times 10^{-1}$ | 0.172 | *** |
| Age<sup>2</sup> | $-3.00 \times 10^{-5}$ | -3.20 | $1.647 \times 10^{-3}$ | $-3.35 \times 10^{-1}$ | $-8.02 \times 10^{-2}$ | 0.054 | ** |
| CSPT (C):Age | $-3.25 \times 10^{-4}$ | -2.56 | $1.088 \times 10^{-2}$ | $-2.16 \times 10^{-1}$ | $-2.90 \times 10^{-2}$ | 0.014 | * |
| ATR (C):Age | $-4.60 \times 10^{-4}$ | -3.70 | $2.456 \times 10^{-4}$ | $-2.65 \times 10^{-1}$ | $-8.11 \times 10^{-2}$ | 0.032 | *** |
| PTR (C):Age | $1.25 \times 10^{-4}$ | 0.98 | $3.266 \times 10^{-1}$ | $-4.74 \times 10^{-2}$ | $1.40 \times 10^{-1}$ | 0.000 | |
| FP (C):Age | $-5.59 \times 10^{-4}$ | -4.35 | $1.709 \times 10^{-5}$ | $-3.06 \times 10^{-1}$ | $-1.16 \times 10^{-1}$ | 0.045 | *** |
| CSPT (C):Age<sup>2</sup> | $3.11 \times 10^{-6}$ | 0.42 | $6.721 \times 10^{-1}$ | $-7.84 \times 10^{-2}$ | $1.21 \times 10^{-1}$ | 0.000 | |
| ATR (C):Age<sup>2</sup> | $-5.82 \times 10^{-6}$ | -0.82 | $4.144 \times 10^{-1}$ | $-1.37 \times 10^{-1}$ | $5.66 \times 10^{-2}$ | 0.000 | |
| PTR (C):Age<sup>2</sup> | $7.73 \times 10^{-6}$ | 1.05 | $2.933 \times 10^{-1}$ | $-4.64 \times 10^{-2}$ | $1.53 \times 10^{-1}$ | 0.000 | |
| FP (C):Age<sup>2</sup> | $6.45 \times 10^{-6}$ | 0.87 | $3.839 \times 10^{-1}$ | $-5.60 \times 10^{-2}$ | $1.45 \times 10^{-1}$ | 0.000 | |
| Sex | $-1.28 \times 10^{-2}$ | -2.33 | $2.184 \times 10^{-2}$ | $-2.34 \times 10^{-1}$ | $-1.99 \times 10^{-2}$ | 0.042 | * |

Note: Significance levels: \* $p < 0.05$ ; \*\* $p < 0.01$ , and \*\*\* $p < 0.001$ . Abbreviations: ATR - anterior thalamic radiation, C – combined, CSPT - cortico-striato-pallido-thalamic, FP – frontoparietal, PTR - posterior thalamic radiation.

**Table 5.** Age effect comparisons of mean orientation dispersion index within specific tracts relative to the whole tractogram.

| IV | *b* | *t* | *p* | 95% CI | | $R^2_{Adj.}$ | Sig. |
| --- | --- | --- | --- | --- | --- | --- | --- |
|  |  |  |  | LL | UL |  |  |
| (Intercept) | $2.99 \times 10^{-1}$ | 148.78 | $2.431 \times 10^{-305}$ | 1.43 | 1.77 | 0.985 | *** |
| CSPT (C) | $-2.59 \times 10^{-2}$ | -11.49 | $1.272 \times 10^{-27}$ | -1.30 | $-9.17 \times 10^{-1}$ | 0.186 | *** |
| FP (L) | $-3.20 \times 10^{-2}$ | -14.12 | $4.726 \times 10^{-39}$ | -1.56 | -1.18 | 0.257 | *** |
| FP (R) | $-1.50 \times 10^{-2}$ | -6.61 | $8.567 \times 10^{-11}$ | $-8.37 \times 10^{-1}$ | $-4.53 \times 10^{-1}$ | 0.069 | *** |
| ATR (L) | $-4.83 \times 10^{-2}$ | -21.54 | $1.053 \times 10^{-75}$ | -2.26 | -1.88 | 0.448 | *** |
| ATR (R) | $-4.55 \times 10^{-2}$ | -20.16 | $1.125 \times 10^{-68}$ | -2.14 | -1.76 | 0.415 | *** |
| PTR (C) | $-4.49 \times 10^{-2}$ | -19.97 | $1.132 \times 10^{-67}$ | -2.11 | -1.73 | 0.410 | *** |
| Age | $-4.26 \times 10^{-4}$ | -5.50 | $7.367 \times 10^{-8}$ | $-4.50 \times 10^{-1}$ | $-2.11 \times 10^{-1}$ | 0.078 | *** |
| Age<sup>2</sup> | $-1.93 \times 10^{-5}$ | -4.43 | $1.271 \times 10^{-5}$ | $-4.04 \times 10^{-1}$ | $-1.56 \times 10^{-1}$ | 0.052 | *** |
| CSPT (C):Age | $-1.28 \times 10^{-4}$ | -1.48 | $1.406 \times 10^{-1}$ | $-2.33 \times 10^{-1}$ | $3.32 \times 10^{-2}$ | 0.002 | |
| ATR (L):Age | $1.83 \times 10^{-5}$ | 0.21 | $8.324 \times 10^{-1}$ | $-1.20 \times 10^{-1}$ | $1.46 \times 10^{-1}$ | 0.000 | |
| ATR (R):Age | $-7.80 \times 10^{-5}$ | -0.90 | $3.678 \times 10^{-1}$ | $-1.95 \times 10^{-1}$ | $7.16 \times 10^{-2}$ | 0.000 | |
| PTR (C):Age | $-2.10 \times 10^{-4}$ | -2.43 | $1.550 \times 10^{-2}$ | $-2.97 \times 10^{-1}$ | $-3.04 \times 10^{-2}$ | 0.008 | * |
| FP (L):Age | $1.20 \times 10^{-4}$ | 1.39 | $1.655 \times 10^{-1}$ | $-3.85 \times 10^{-2}$ | $2.28 \times 10^{-1}$ | 0.002 | |
| FP (R):Age | $3.93 \times 10^{-5}$ | 0.45 | $6.511 \times 10^{-1}$ | $-1.01 \times 10^{-1}$ | $1.66 \times 10^{-1}$ | 0.000 | |
| CSPT (C):Age<sup>2</sup> | $-2.15 \times 10^{-6}$ | -0.43 | $6.656 \times 10^{-1}$ | $-1.73 \times 10^{-1}$ | $1.10 \times 10^{-1}$ | 0.000 | |
| ATR (L):Age<sup>2</sup> | $3.87 \times 10^{-6}$ | 0.79 | $4.319 \times 10^{-1}$ | $-8.42 \times 10^{-2}$ | $1.97 \times 10^{-1}$ | 0.000 | |
| ATR (R):Age<sup>2</sup> | $4.74 \times 10^{-7}$ | 0.1 | $9.241 \times 10^{-1}$ | $-1.35 \times 10^{-1}$ | $1.49 \times 10^{-1}$ | 0.000 | |
| PTR (C):Age<sup>2</sup> | $-7.01 \times 10^{-6}$ | -1.41 | $1.591 \times 10^{-1}$ | $-2.43 \times 10^{-1}$ | $4.00 \times 10^{-2}$ | 0.002 | |
| FP (L):Age<sup>2</sup> | $-5.10 \times 10^{-8}$ | -0.01 | $9.919 \times 10^{-1}$ | $-1.43 \times 10^{-1}$ | $1.41 \times 10^{-1}$ | 0.000 | |
| FP (R):Age<sup>2</sup> | $-5.56 \times 10^{-6}$ | -1.11 | $2.664 \times 10^{-1}$ | $-2.23 \times 10^{-1}$ | $6.17 \times 10^{-2}$ | 0.000 | |
| Sex | $4.68 \times 10^{-3}$ | 2.3 | $2.376 \times 10^{-2}$ | $1.41 \times 10^{-2}$ | $1.80 \times 10^{-1}$ | 0.042 | * |

Note. Significance levels: \* $p < 0.05$ ; \*\* $p < 0.01$ , and \*\*\* $p < 0.001$ . Abbreviations: ATR - anterior thalamic radiation, C – combined, CSPT – cortico-striato-pallido-thalamic, FP – frontoparietal, L – left, PTR - posterior thalamic radiation, R – right.

For mean orientation dispersion index, each tract of interest showed lower levels of ODI relative to the whole brain tractogram (*p* < 8.567 × 10^−11^, adjusted *R^2^* ranged from 0.07 to 0.45). The whole brain tractogram evidenced both a significant linear age (*p* = 7.367 × 10^−8^, adjusted *R^2^* = 0.08) and quadratic age effect (*p* = 1.271 × 10^−5^, adjusted *R^2^* = 0.05). For the linear age trajectory, the combined posterior thalamic radiation tract had a steeper slope with age compared to the whole brain tractogram (*p* = 0.016, adjusted *R^2^* = 0.08). The mean ODI or complexity of neurite orientation decreased faster with age for the combined PTR tract compared to the whole brain tractogram. However, for the quadratic age trajectory, there were no significant differences in each of the TOIs relative to the whole brain tractogram suggesting that the other TOIs followed a similar quadratic trajectory. These LME analyses reveal that the average amount of microstructural properties in each tract of interest differed from the whole brain tractogram with more in the case of NDI and less for the case of ODI. For both neurite microstructural properties, these LME analyses confirm that most TOIs follow a quadratic trajectory through the adult lifespan that is similar to the whole brain tractogram. Also, for both metrics, the linear age trajectory was similar for most tracts of interest compared to the whole brain tractogram with some TOIs showing accelerated linear decline with age.

### 3.2. Spatial Gradients of Neurite Microstructural Properties

To determine whether the individual-level variations of spatial gradients are consistent at the group-level, a GLM was performed for each cardinal direction with the spatial gradient estimate as the DV and the IV being the intercept while controlling for the mean-centered spatial autocorrelation. Significant spatial gradients were observed across each direction and microstructural property in most white matter tracts (see **Figure 4** for NDI and **Figure 5** for ODI). For NDI in the medial-to-lateral direction (NDI_ML_), a negative spatial gradient was observed in the bilateral whole brain tractogram, right cortico-striato-pallido-thalamic, bilateral posterior thalamic radiation, and bilateral frontoparietal tracts such that greater neurite density was observed in the medial portion relative to the lateral portion. For NDI in the posterior-to-anterior direction (NDI_PA_), a negative spatial gradient was observed for the bilateral whole brain tractogram such that more neurite density was observed in the posterior portion relative to the anterior portion. Additionally, for NDI_PA_, a positive spatial gradient was observed in the bilateral cortico-striato-pallido-thalamic, bilateral posterior thalamic radiation, and bilateral frontoparietal tracts such that more neurite density was observed in the anterior portion relative to the posterior portion. For NDI in the ventral-to-dorsal direction (NDI_VD_), a negative spatial gradient was observed in all tracts of interest such that more neurite density was observed in the ventral portion relative to the dorsal portion.

**Figure 4.**
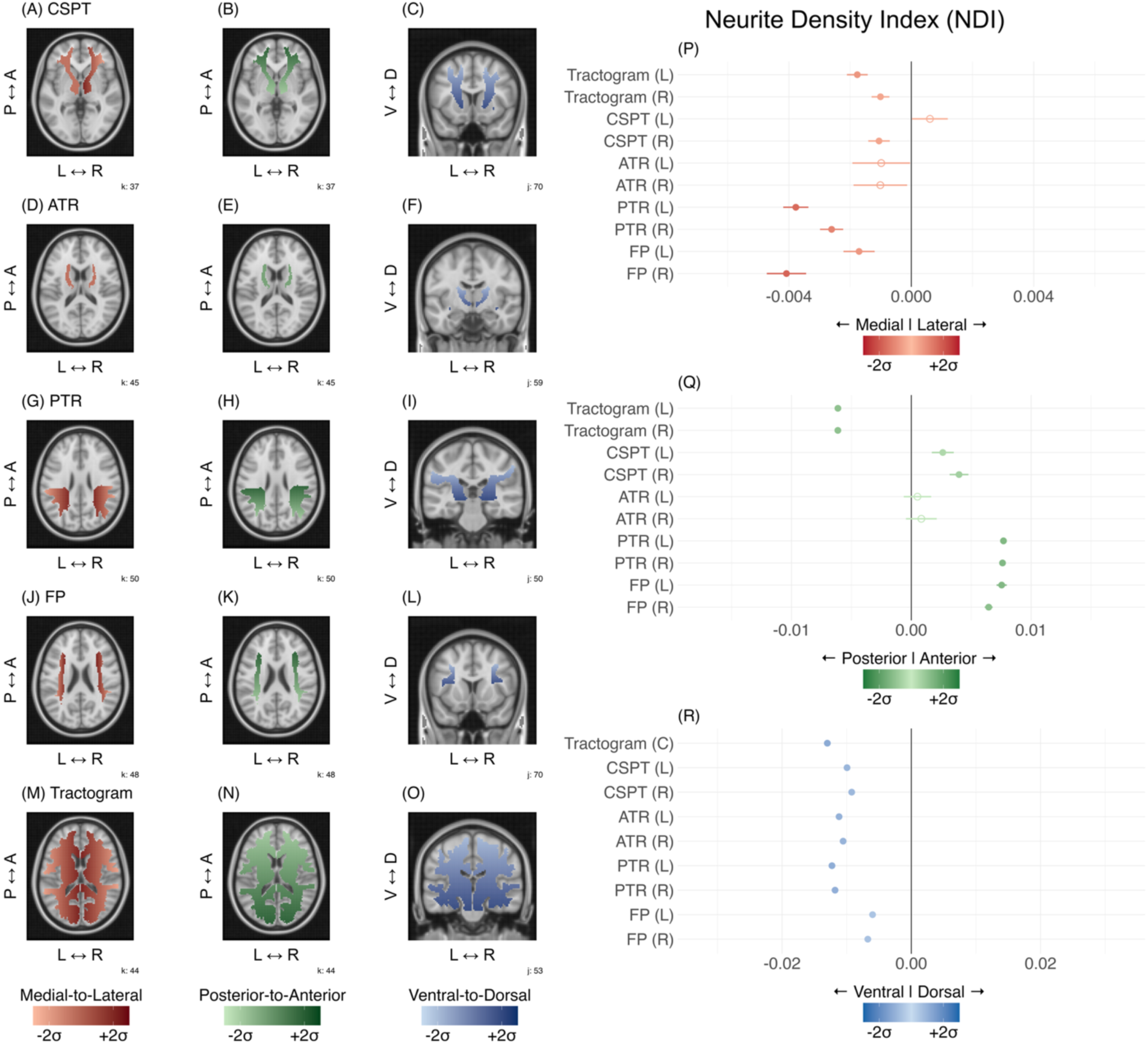
Spatial gradients of neurite density index for each tract of interest along the medial-to-lateral, posterior-to-anterior, and ventral-to-dorsal directions. Note: Non-significant spatial gradients are represented as a solid color (A-O) and as unfilled circles (P-R). Error bars represent the 95% confidence interval. Abbreviations: ATR – anterior thalamic radiations, C – combined, CSPT - cortico-striato-pallido-thalamic tract, FP – frontoparietal tract, L – left, PTR – posterior thalamic radiations, R - right.

**Figure 5.**
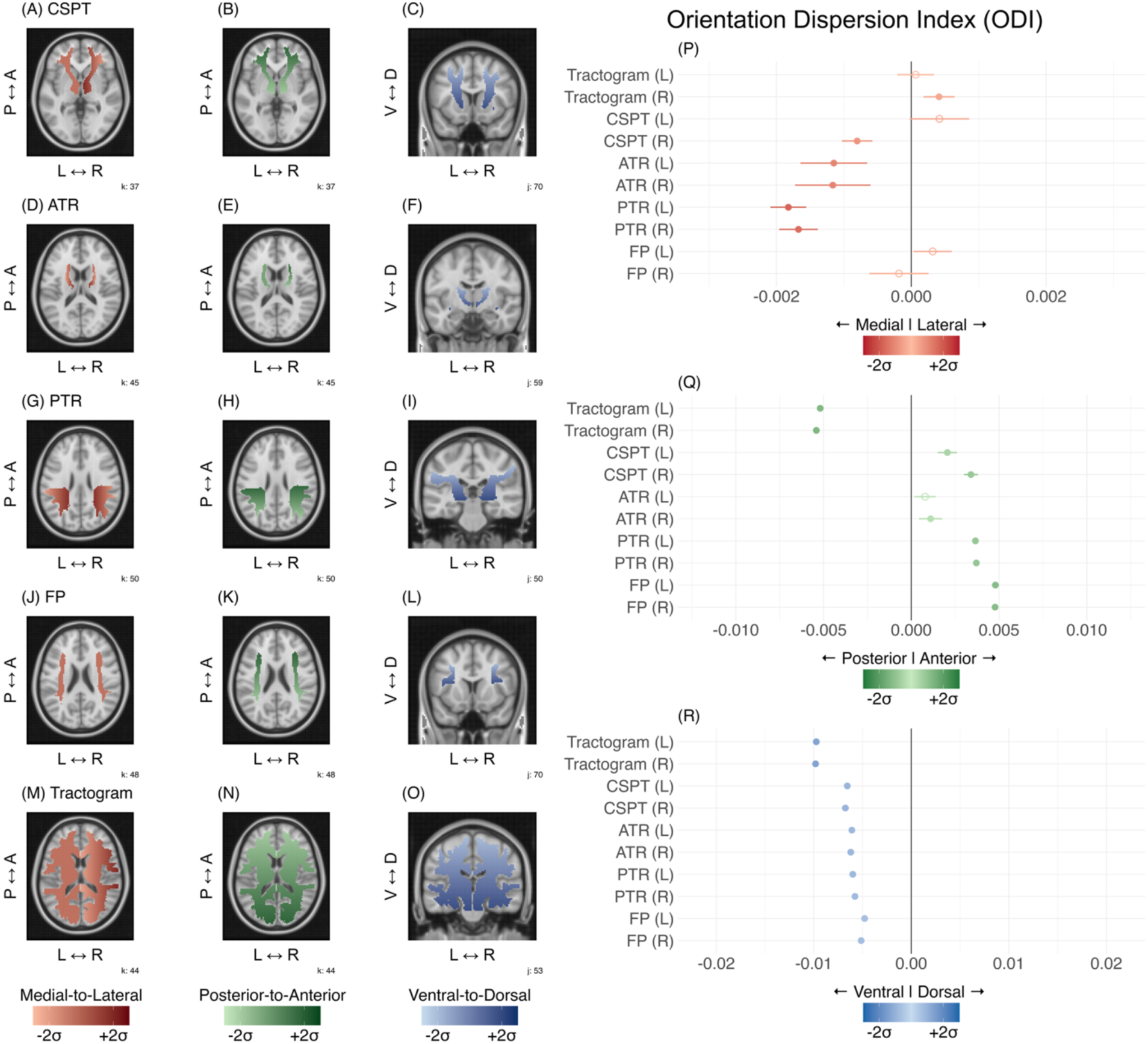
Spatial gradients of orientation dispersion index for each tract of interest along the medial-to-lateral, posterior-to-anterior, and ventral-to-dorsal direction. Note: Non-significant spatial gradients are represented as a solid color (A-O) and as unfilled circles (P-R). Error bars represent the 95% confidence interval. Abbreviations: ATR – anterior thalamic radiations, C – combined, CSPT - cortico-striato-pallido-thalamic tract, FP – frontoparietal tract, L – left, PTR – posterior thalamic radiations, R - right.

For ODI (**Figure 5**) in the medial-to-lateral direction (ODI_ML_), a negative spatial gradient was observed in the right CSPT as well as bilaterally in the ATR and PTR tracts such that more dispersion of neurite orientation was observed in the medial portion relative to the lateral portion. Additionally, for ODI_ML_, a positive spatial gradient was observed for the right whole brain tractogram such that more dispersion of neurite orientation was observed for the lateral relative to the medial portion. For ODI in the posterior-to-anterior direction (ODI_PA_), a significant negative spatial gradient was observed in bilateral whole brain tractogram such that more dispersion of neurite orientation was observed in the posterior portion relative to the anterior portion. Additionally, for ODI_PA_, a significant positive spatial gradient was observed in the right ATR as well as bilaterally in the CSPT, PTR, and FP tracts such that more dispersion of neurite orientation was observed in the anterior portion relative to the posterior portion. For ODI in the ventral-to-dorsal direction (ODI_VD_), a significant negative spatial gradient was observed bilaterally in all tracts of interest such that more dispersion of neurite orientation was observed in the ventral portion relative to the dorsal portion. For most TOIs, individually-estimated spatial gradients were consistent and reliable at the group-level across both microstructural properties and each cardinal direction.

### 3.3. Age Effects on Spatial Gradients of Neurite Microstructural Properties

To examine the effects of age on these spatial gradients of microstructural properties, a GLM was performed with the spatial gradient estimate as the DV with linear and quadratic age as the IVs while controlling for sex and spatial autocorrelation for each microstructural property, cardinal direction, and TOI. Significant effects were found for both linear and quadratic age on the spatial gradients of microstructural properties (see **Table 6** and **Figure 6**). For NDI, significant age effects were observed in the posterior-to-anterior (NDI_PA_) and ventral-to-dorsal (NDI_VD_) directions, but not in the medial-to-lateral direction. For NDI_PA_, the spatial gradient decreased (or became more homogenous between the two ends of the axis) with age in the bilateral tractogram and the left frontoparietal tract. In other words, for these tracts, the amount of NDI in the posterior and anterior portions became more similar with increasing age (*p*’s < 5.362 × 10^−3^, adjusted *R^2^* ranged from 0.07 to 0.10). For NDI_VD_, a quadratic age effect was found for the whole brain tractogram (*p* = 7.569 × 10^−4^, adjusted *R^2^* = 0.10) and a linear age effect was found for the right CSPT (*p* = 3.713 × 10^−6^, adjusted *R^2^*= 0.20). For the whole brain tractogram, the spatial gradient of NDI increased slightly until middle-age and subsequently decreased (or became more homogenous between the two ends of the axis) in older adulthood. In other words, for the whole brain tractogram, more NDI was found in the ventral portion compared to the dorsal portion in younger adulthood, then this discrepancy became steeper with more NDI in the ventral portion until middle adulthood, and then the discrepancy between these portions became less steep in older adulthood. For the right CSPT, the spatial gradient decreased (or became more homogenous) with age. For ODI, a significant age effect was observed for the ventral-to-dorsal direction (ODI_VD_), but not for either the medial-to-lateral or posterior-to-anterior directions. For ODI_VD_, the spatial gradient decreased (or became more homogenous) linearly with age within the right CSPT tract (*p* = 5.962 × 10^−6^, adjusted *R^2^* = 0.19).

**Figure 6.**
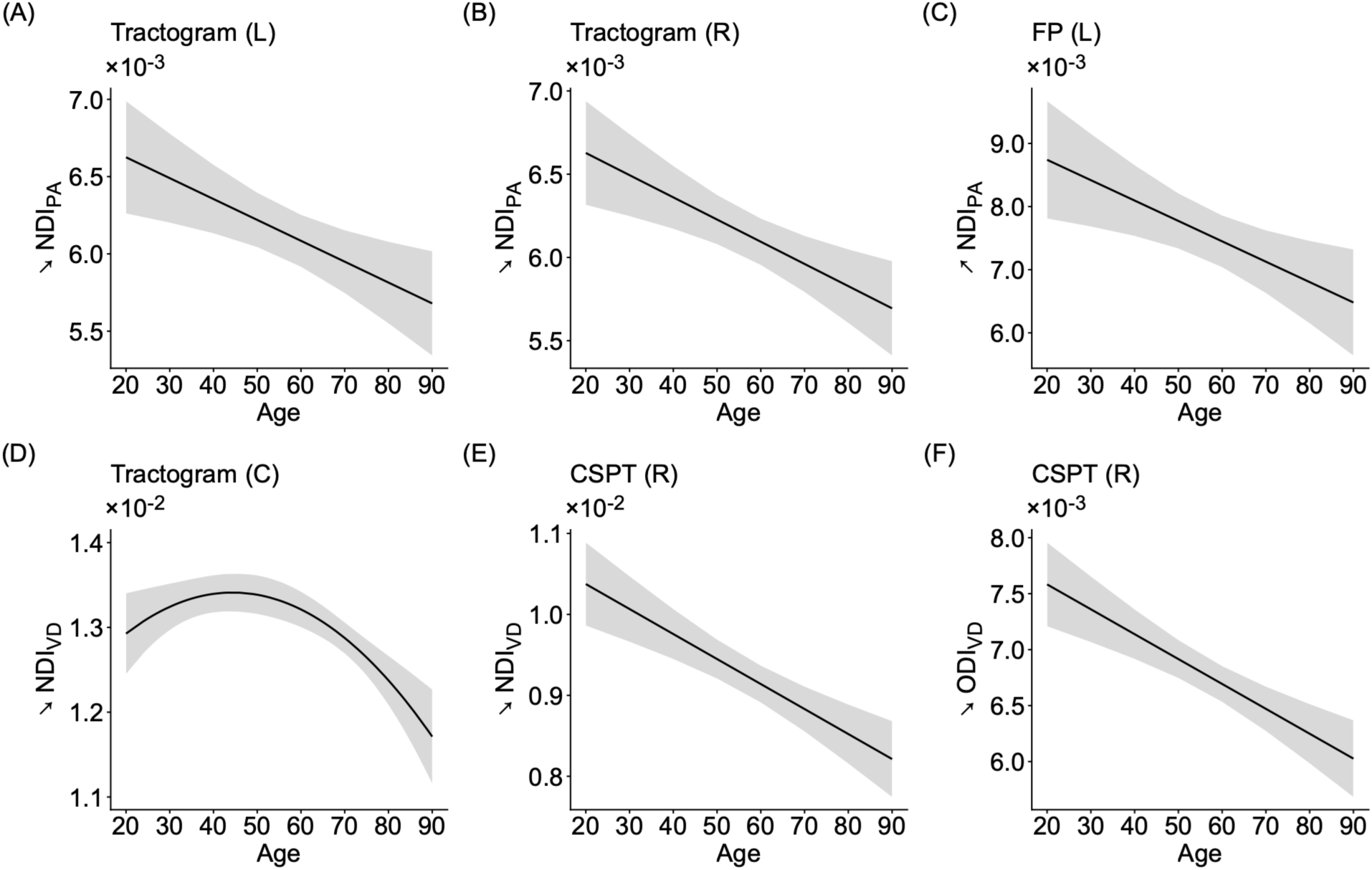
Significant linear and quadratic age effects on spatial gradients of neurite microstructural properties. Spatial gradient in the posterior-to-anterior weakened (became more homogenous) with age for the bilateral tractogram and left frontoparietal tract (A-C). Spatial gradient in the ventral-to-dorsal direction strengthened towards middle adulthood and then weakened in older adulthood within the whole brain tractogram (D). Spatial gradient in the ventral-to-dorsal direction weakened (became more homogenous) with age in the bilateral cortico-striato-pallido-thalamic tract for NDI (E) and ODI (F). Note: The y-axis represents the estimated spatial gradient in its respective direction when controlling for sex. The bands represent the 95% confidence interval. Abbreviations: C – combined, CSPT - cortico-striato-pallido-thalamic tract, FP – frontoparietal tract, L – left, R - right.

**Table 6.** Linear and quadratic age effects for spatial gradients of microstructural properties.

| Metric | IV | Dir. | TOI | $b$ | $t$ | $p$ | 95% CI | | $R^2_{Adj.}$ |
| --- | --- | --- | --- | --- | --- | --- | --- | --- | --- |
|  |  |  |  |  |  |  | LL | UL |  |
| NDI<sub>PA</sub> | Age | ↘ | Tractogram (L) | $-1.35 \times 10^{-5}$ | -3.042 | $3.033 \times 10^{-3}$ | $-2.23 \times 10^{-5}$ | $-4.69 \times 10^{-6}$ | 0.078 |
| | | ↘ | Tractogram (R) | $-1.33 \times 10^{-5}$ | -3.513 | $6.674 \times 10^{-4}$ | $-2.09 \times 10^{-5}$ | $-5.80 \times 10^{-6}$ | 0.101 |
| | | ↗ | FP (L) | $-3.23 \times 10^{-5}$ | -2.854 | $5.362 \times 10^{-3}$ | $-5.48 \times 10^{-5}$ | $-9.81 \times 10^{-6}$ | 0.073 |
| NDI<sub>VD</sub> | Age<sup>2</sup> | ↘ | Tractogram (C) | $-8.16 \times 10^{-7}$ | -3.476 | $7.569 \times 10^{-4}$ | $-1.28 \times 10^{-6}$ | $-3.50 \times 10^{-7}$ | 0.100 |
| NDI<sub>VD</sub> | Age | ↘ | CSPT (R) | $-3.09 \times 10^{-5}$ | -4.924 | $3.713 \times 10^{-6}$ | $-4.33 \times 10^{-5}$ | $-1.84 \times 10^{-5}$ | 0.200 |
| ODI<sub>VD</sub> | Age | ↘ | CSPT (R) | $-2.22 \times 10^{-5}$ | -4.800 | $5.962 \times 10^{-6}$ | $-3.14 \times 10^{-5}$ | $-1.30 \times 10^{-5}$ | 0.188 |

*Note:* Direction represents the spatial gradient direction for that particular tract, microstructural property, and cardinal direction. Abbreviations: C – combined, CSPT – cortico-striato-pallido-thalamic, Dir. – direction, FP – frontoparietal, L – left, NDI – Neurite Density Index, ODI – Orientation Dispersion Index, PA – posterior-to-anterior, R – right, VD – ventral-to-dorsal.

For spatial gradients of both neurite microstructural properties, most tracts and cardinal directions showing significant age effects followed a linear age trajectory with the spatial gradient becoming more homogenous with increasing age. However, one significant result (**Figure 6D**) revealed a non-linear, quadratic relationship with age such that spatial gradient increased slightly until middle-age and then became more homogenous in older adulthood.

To determine how age influenced microstructural properties at each slice (or spatial patterns of age effects) along a cardinal direction similar to prior studies using DTI (Hoagey et al., 2019), a GLM was performed with mean microstructural property for a particular slice as the DV with both linear and quadratic age as the IVs while controlling for sex. For both neurite microstructural properties, an omnibus or combined effect of age effect was found on nearly all slices for each cardinal direction (See **Figure 7**). For NDI, the combined age effect size peaked near the midway point along each cardinal direction with the maxima occurring within the lateral, posterior, and dorsal portions. For ODI, the combined age effect size also peaked near the midway point along each cardinal direction with the maxima occurring within in the medial and dorsal portions. For ODI in the posterior-to-anterior direction, the peaks were essentially on the midway point. Both neurite microstructural properties exhibited a non-linear profile of the combined age effect size (or age gradient) along the slices of each cardinal direction with peaks occurring near the midpoint of each axis.

**Figure 7.**
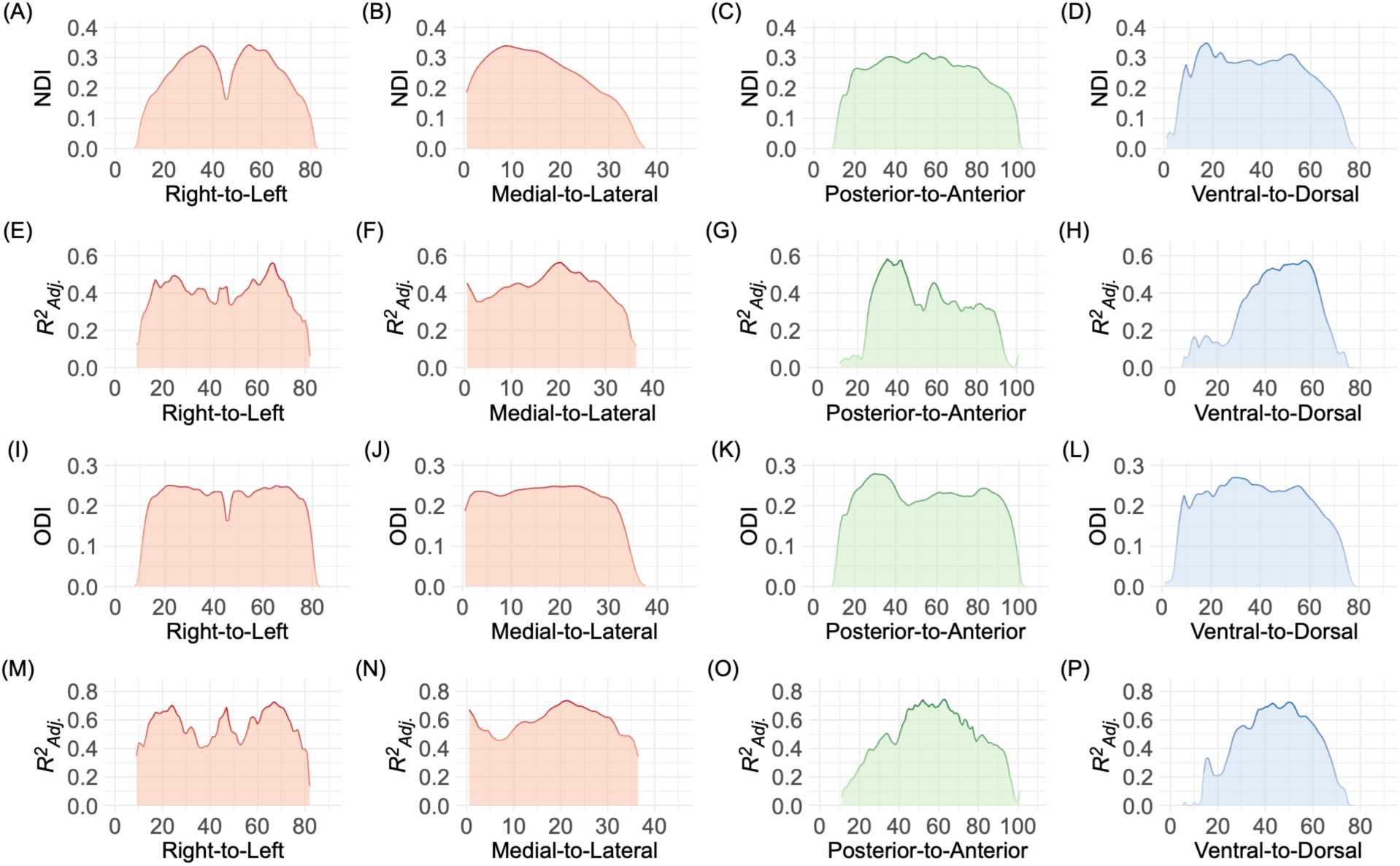
Mean neurite microstructural property across subjects (A-D; I-L) and its associated age effect size (E-H; M-P) for each slice in the right-to-left, medial-to-lateral, posterior-to-anterior, and ventral-to-dorsal directions in the whole brain tractogram. For NDI, there is an age-effect pattern along the cardinal axes with larger age effects generally in the lateral, posterior, and dorsal portions. For ODI, there is an age effect pattern along the cardinal axes with larger age effects generally in the medial and dorsal portion. Interestingly, the effect size tended to peak midway along the cardinal axes. The effect size is the adjusted *R^2^* of the combined effect of linear and quadratic age. Abbreviations: NDI – neurite density index, ODI – orientation dispersion index

## 4. DISCUSSION

The current study investigated how neurite microstructural properties in white matter structural connections related to the cortical-striato-pallido-thalamic loop were associated with age across the adult lifespan. For NDI and ODI in these white matter tracts, both average neurite microstructure and spatial gradients of neurite microstructure along the cardinal axes were demonstrated to be detrimentally influenced by age and differed by tract and gradient direction.

### 4.1. Age Effects on Average Neurite Microstructural Properties

Average neurite density and orientation dispersion indices, which are proxies respectively for axonal amount in white matter tissue and axonal complexity, mostly followed an inverted-U quadratic relationship with age, increasing towards middle adulthood and then decreasing in older adulthood. These findings are consistent with prior studies of neurite microstructural properties that also showed either this nonlinear relationship with age or a linear decline with age in major white matter bundles (Billiett et al., 2015; Cox et al., 2016; Gozdas et al., 2021; Lawrence et al., 2021). These findings are also consistent with other studies using different microstructural properties such as FA and myelin water fraction revealing an inverted-U quadratic trajectory with age and MD and myelin sheath thickness (using g-ratio) having a U-shaped quadratic relationship with age (Beck et al., 2021; Bouhrara et al., 2021; Cox et al., 2016; Kochunov et al., 2021; Webb et al., 2020a; Qian et al., 2020). Compared to the whole white matter tractogram, some tracts and metrics showed age-associations on par with the tractogram age effects, whereas some tracts and metrics showed aging effects beyond that of the general tractogram. For axonal density, the CSPT, ATR, and FP tracts had a steeper age slope than the tractogram, with the frontoparietal tract evidencing the steepest effect (followed by ATR, and then CSPT). For axonal complexity, the PTR tract demonstrated a steeper age slope compared to the white matter tractogram. This pattern of differential aging is interesting both in its regional pattern but also in the type of aging process evidenced – more white matter seems to naturally decrease in neurite density with age, but only the posterior thalamic radiations had an accelerated aging pattern in orientation dispersion of axon bundles.

### 4.2. Spatial Gradients of Neurite Microstructural Properties

Significant spatial gradients were found for each cardinal direction and for each neurite microstructural property. These findings align with prior studies investigating spatial gradients in structural white matter connections. For corticostriatal connections, more anterior portions of the striatum are more connected with anterior and associative areas of the cortex (Jarbo & Versytnen, 2015; Versytnen et al., 2012). For thalamocortical connections, more medial portions of the thalamus are connected with more medial and anterior portions of the cortex (Mengxing et al., 2023). Our spatial gradients of white matter connections related to the cortico-striato-pallido-thalamic loop also appear to follow a similar pattern of more axonal properties (greater density and dispersion) in the anterior and medial portions of the tracts. Interestingly, all tracts of interest showed a reliable spatial gradient in the ventral-to-dorsal direction with more axonal density and dispersion in the ventral portions. Given the consistency of this spatial gradient across tracts of varying location and shape, this ventral-to-dorsal spatial gradient may be a ubiquitous pattern of axons in the brain. Although the ubiquity of the pattern needs to be confirmed with additional tracts, the greater axonal properties in the ventral portion may be due to phylogenic conservation that may be reflected in gray matter. In gray matter tissue, the ventral portion has less evolutionary and developmental expansion relative to the dorsal portions (Hill et al., 2010).

### 4.3. Age Effects on Spatial Gradients of Neurite Microstructural Properties

Interestingly, these neurite spatial gradients were age-sensitive. Axonal density gradient in the posterior-to-anterior direction became more homogenous with increasing age in the bilateral white matter tractogram and the left frontoparietal tract (i.e., the gradient pattern was lost with age). Axonal density in the ventral-to-dorsal direction followed a nonlinear, inverted-U with age in the whole brain tractogram and linear decline with age in the right CSPT tract, where these became more homogenous with increasing age. Given that the striato-pallido-thalamic projections have been shown to be segregated by striatal subcompartments such as the striosome and matrix (Funk et al., 2023), we speculate that the microstructural properties in striatal subcompartments itself may also become more homogeneous with age and this may occur in tandem with the increasing homogeneity of these tracts with age. This aging pattern resembles the dedifferentiation found in large functional networks with increasing age (Malagurki et al., 2020; Chan et al., 2014). Interestingly, in younger adults striosome-like and matrix-like subcompartments within the striatum revealed differences in resting-state functional connectivity across networks (Sadiq et al., 2025); however, dedifferentiation of functional connections with these striatal subcompartments with age remains to be examined. It may be a general aging property of the brain that homogeneity or dedifferentiation increases, as it has been observed in white matter tracts, gray matter tissue, and functional connections.

### 4.4. Limitations and Future Directions

These results should be interpreted in the context of their caveats and limitations. First, spatial gradients may be more complex than captured in the current analyses and more complex models may be needed for further examination. For example, neurite microstructural properties may be quadratic with higher amounts towards the center of the region. Future studies should model and quantitatively compare which level of polynomial models of spatial gradients improve estimation and to what degree of nonlinearity. Second, white matter hyperintensities, representing lesions that increase in volume and frequency with age (Hoagey et al., 2021; Raz et al., 2012) have been shown to be associated with decreased NDI (James et al., 2023; Raghaven et al., 2021), decreased FA, decreased myelin, and increased MD (Hoagey et al., 2021; James et al., 2023; Raghavan et al., 2021). Given the association of white matter hyperintensities on microstructural properties, examination of this as a potential confound (or potential factor of interest) may be warranted for future studies. Third, relatively large and long tracts were selected for the present study given the current spatial resolution. However, tractography of smaller segments of this loop such as the striatopallido and pallidothalamic tracts would allow for a finer granularity examination of neurite microstructural properties and their gradients. Thus, future studies should examine the specificity of these findings in more granular tract segments. Fourth, partial volume effects (PVE) have been associated with DTI microstructural properties related to bundle size, bundle curvature, and bundle alignment with the acquisition grid, and specifically PVE effects was stronger for smaller bundles than larger ones, PVE was nonlinear for curvature and depended upon metric, and that PVE was stronger when bundles were not in alignment with the acquisition grid (Vos et al., 2020). While less likely to have been an issue for the current multi-shell study with longer tracts, future studies should examine how tract volume, curvature, and proportion of tract within a voxel may influence these neurite microstructural properties, especially when measuring spatial gradients. Another important future study comes from the speculation of the relationship between the spatial gradients of gray matter tissue. Specifically, we speculated that spatial gradients of white matter structural connections are becoming more homogenous in tandem or following the homogeneity of spatial gradients of gray matter regions. Future studies should explicitly examine whether the spatial gradients of these two tissue types covary together or represent separate and unique properties with age. If the spatial gradients of these two tissue types are confirmed to covary together, future studies should also determine their temporal order in longitudinal studies, as cross-sectional age studies are limited by cohort effects and cannot speak to within-person changes over time.

### 4.4. Conclusion

The current study found that neurite microstructural properties in white matter structural connections related to the cortico-striato-pallido-thalamic loop were differentially sensitive to the effects of age. Average neurite microstructural properties of these structural connections followed mostly an inverted-U quadratic relationship with age, while homogeneity of the spatial gradients mostly followed a linear relationship with age. This compromise of the tract bundle architecture may occur in tandem with or be due to the despecialization of regional subcompartments found in gray matter tissue such as the striatal subcompartments. The present study adds to the current literature by demonstrating various spatial gradients of neurite microstructural properties along the white matter tracts of an important fundamental cortico-subcortical loop that are age-sensitive and differential in their age trajectories.

## Acknowledgements

This work was supported, in part, by grants from the National Institute of Health (R01AG056535 and R01AG057537).

